# Challenges in the discovery of tumor-specific alternative splicing-derived cell-surface antigens in glioma

**DOI:** 10.1101/2023.10.26.564156

**Authors:** Takahide Nejo, Lin Wang, Kevin K. Leung, Albert Wang, Senthilnath Lakshmanachetty, Marco Gallus, Darwin W. Kwok, Chibo Hong, Lee H. Chen, Diego A. Carrera, Michael Y. Zhang, Nicholas O. Stevers, Gabriella C. Maldonado, Akane Yamamichi, Payal Watchmaker, Akul Naik, Anny Shai, Joanna J. Phillips, Susan M. Chang, Arun P. Wiita, James A. Wells, Joseph F. Costello, Aaron A. Diaz, Hideho Okada

**Affiliations:** Department of Neurological Surgery, University of California, San Francisco, CA, USA; Department of Pharmaceutical Chemistry, University of California, San Francisco, CA, USA; Department of Laboratory Medicine, University of California, San Francisco, CA, USA; Helen Diller Family Comprehensive Cancer Center, University of California, San Francisco, CA, USA; Department of Pathology, University of California, San Francisco, CA, USA; Department of Bioengineering and Therapeutic Sciences, University of California, San Francisco, CA, USA; Chan Zuckerberg Biohub, CA, USA; The Parker Institute for Cancer Immunotherapy, CA, USA; Department of Cellular and Molecular Pharmacology, University of California, San Francisco, CA, USA

**Keywords:** Glioma, Alternative splicing, Neoantigen, Neojunction, Bulk RNA-sequencing, Long-read sequencing, Intratumoral heterogeneity, Proteomics

## Abstract

**Background:** Despite advancements in cancer immunotherapy, solid tumors remain formidable challenges. In glioma, profound inter-and intra-tumoral heterogeneity of antigen landscape hampers therapeutic development. Therefore, it is critical to consider alternative sources to expand the repertoire of targetable (neo-)antigens and improve therapeutic outcomes. Accumulating evidence suggests that tumor-specific alternative splicing (AS) could be an untapped reservoir of neoantigens.

**Results:** In this study, we investigated tumor-specific AS events in glioma, focusing on those predicted to generate major histocompatibility complex (MHC)-presentation-independent, cell-surface neoantigens that could be targeted by antibodies and chimeric antigen receptor (CAR)-T cells. We systematically analyzed bulk RNA-sequencing datasets comparing 429 tumor samples (from The Cancer Genome Atlas [TCGA]) and 9,166 normal tissue samples (from the Genotype-Tissue Expression project [GTEx]), and identified 13 AS events in 7 genes predicted to be expressed in more than 10% of the patients, including *PTPRZ1* and *BCAN*, which were corroborated by an external RNA-sequencing dataset. Subsequently, we validated our predictions and elucidated the complexity of the isoforms using full-length transcript amplicon sequencing on patient-derived glioblastoma cells. However, analyses of the RNA-sequencing datasets of spatially mapped and longitudinally collected clinical tumor samples unveiled remarkable spatiotemporal heterogeneity of the candidate AS events. Furthermore, proteomics analysis did not reveal any peptide spectra matching the putative neoantigens.

**Conclusions:** Our investigation illustrated the diverse characteristics of the tumor-specific AS events and the challenges of antigen exploration due to their notable spatiotemporal heterogeneity and elusive nature at the protein levels. Redirecting future efforts toward intracellular, MHC-presented antigens could offer a more viable avenue.

## Background

Over the last decade, cancer immunotherapy, including immune checkpoint blockade (ICB) and chimeric antigen receptor (CAR)-T cell therapy, has revolutionized treatments for various types of cancers, notably encompassing hematological malignancies and melanoma [1–3]. However, many patients, particularly those with solid tumors, including glioma, have not yet experienced commensurate therapeutic benefits [4–6].

The sparsity of targetable antigens is a crucial obstacle for immunotherapeutic approaches targeting specific antigens, such as CAR-T, genetically engineered T-cell-receptor (TCR)-T cell, and vaccines [7,8]. For the development of such therapeutics, multiple critical factors must be considered. For safety and efficacy, it is crucial that these targets are both robustly expressed and exclusively present on tumor cells, and absent on non-tumor cells and healthy tissues. In particular, antigen tumor-specificity must be assured to avoid on-target/off-tumor cross-reactions of the infused or induced immune cells, which can sometimes be lethal [9–11]. Therefore, tumor-specific antigens (neoantigens) arising from somatic mutations have emerged as attractive targets [12]. In particular, neoantigens shared among patients would be more suitable for an off-the-shelf approach; however, the number of such neoantigens currently available is limited. For example, in the case of glioma, only a few shared neoantigens have been identified and investigated in clinical trials [13]. These include mutations, such as *IDH1* R132H and *EGFR*vIII in adult glioma [14,15] and H3.3 K27M in pediatric glioma [16,17]. Additionally, spatio-temporal and inter-and intra-patient heterogeneity amplify this challenge, as glioma is known for its remarkable heterogeneity [18]. It is anticipated that the single neoantigen-oriented therapies are likely to fail in heterogenous tumors due to tumor clonal evolution and antigen loss [15,19]. As such, it is crucial to expand the repertoire of antigens so that more patients who are otherwise ineligible for currently available therapeutic options can benefit, increasing the chances of treatment success.

Recent studies have shown that neoantigens can arise from sources outside of somatic mutations, such as aberrant RNA splicing specific to cancers [7,8]. Two studies analyzing the RNA sequencing (RNA-seq) data from The Cancer Genome Atlas (TCGA) revealed that tumor-specific alternative splicing (AS) events, or “neojunctions,” are abundant and may contribute to expanding the repertoires of therapeutic targets for cancer immunotherapy [20,21]. Kahles et al. also showed that *IDH1* mutations, observed in approximately 24% of adult glioma cases [22], are associated with increased genome-wide splicing disruption [20]. The studies collectively reported that some tumor-specific AS events are shared among the patients, implicating the translational possibility of cancer immunotherapy targeting these AS-derived neoantigens. Furthermore, other recent studies have provided proof-of-concept data showing that certain AS-derived neoepitopes exhibit protein-level expression and demonstrate immunogenicity [23–25]. Altogether, these observations indicate that exploring cancer-specific AS may expand the repertoire of neoantigens, particularly in glioma, contributing to the development of successful cancer immunotherapy.

In this study, we investigated tumor-specific AS events in glioma as a potential source of untapped neoantigens, with a particular focus on putative cell-surface antigen candidates derived from these events. Importantly, because major histocompatibility complex (MHC)-epitope-type neoantigens have been studied using different analytical approaches and reported separately [26,27], our investigation exclusively focused on the AS events occurring at the cell-surface that are independent of MHC presentation. These cell-surface candidates could serve as targets for antibodies and CAR-T cells in future therapeutic development. Our *in silico* analyses revealed encouraging findings. However, the examination of our unique datasets obtained from clinical tumor samples, including spatially mapped and longitudinally collected specimens, unveiled the unprecedentedly diverse landscape of the candidate tumor-specific AS events. These investigations, along with proteomics data, have shed light on the challenges associated with this discovery strategy.

## Results

### Screening analyses to identify the tumor-specific AS events as potential sources for cell-surface neoantigens

For screening purposes, we developed a discovery analysis workflow with a concept similar to the study of Kahles *et al*.[20]. However, we applied more stringent criteria and focused on avoiding on-target-off-tumor effects in future therapeutic development (**Fig. 1a**). We compared bulk RNA-seq data of adult diffuse glioma in TCGA datasets [28] with those of all normal tissues in Genotype-Tissue Expression project (GTEx) [29]. First, we identified 429 out of 663 primary tumor cases in study projects TCGA-GBM and -LGG, which met our criteria of ABSOLUTE-estimated tumor purity ≥ 60% (median, 82%; range, 60–100%), for subsequent analyses (**Supplementary Table S1**) [30]. The pass-filter TCGA-glioma cohort was composed of 166 glioma/glioblastoma, IDH-wildtype (IDHwt), 140 astrocytoma, IDH-mutant, 1p/19q-non-codeleted (IDH-A), and 123 oligodendroglioma, IDH-mutant, 1p/19q-codeleted (IDH-O) cases [31]. Simultaneously, we obtained bulk RNA-seq data of 9,166 human normal tissues collected from various organs throughout the bodies, including 1,436 brain samples, from the GTEx dataset (**Supplementary Table S2**) [29].

Our analysis workflow initially detected a total of 366,734 distinct splicing junctions with ≥ 10 supportive junction read counts in chromosomes 1–22, X, and Y across 429 tumor samples in the TCGA-glioma dataset (**Fig. 1b**). Among these, 168,286 events were defined as non-annotated based on the absence in the Ensembl reference gene-transfer format (GTF) file. Of these, 9,041 events were found within the genomic regions of the “surfaceome” genes with an average expression of transcript-per-million (TPM) ≥ 10 in the TCGA-glioma dataset. Emphasizing tumor-specificity, we carefully compared the analysis results between TCGA-glioma and GTEx, and retained only events with a positive sample rate (the number of event-positive samples divided by the total number of samples) < 1% in GTEx samples, which resulted in 5,947 “tumor-specific” events passing the filter. Finally, aiming for an off-the-shelf approach in future therapeutic applications, we defined events as “shared” if they were commonly found in ≥ 10% cases in any of the disease groups (IDHwt, IDH-A, or IDH-O) within the TCGA-glioma dataset, resulting in 66 events retained. Notably, an exon skipping event of exons 2–7 in the *EGFR* gene was identified within the list, exclusively in the IDHwt group of the analysis cohort. The event was consistent with the genomic rearrangement mutation *EGFR*vIII, relatively common (∼20%) in IDH-wildtype glioblastoma [32]. *EGFR*vIII is not an alternative splicing event but a genomic rearrangement with an in-frame deletion of exons 2–7, affecting the extracellular domains but retaining the intact transmembrane and intracellular components, leading to constitutive activation of downstream pathways, such as the PI3K/Akt signaling [32]. We utilized *EGFR*vIII as a positive control throughout the downstream analyses hereafter.

Furthermore, we investigated intron-retention (IR) type events separately using IRFinder [33]. Similar to the exon-exon junction investigation, we compared the intron expression profiles of tumor samples from IDHwt (n = 166), IDH-A (n = 140), and IDH-O (n = 123) in the TCGA-glioma dataset with those from normal brain samples in the GTEx dataset (n = 1,436). Through differential intron retention analysis, we identified 893 distinct events that exhibited significantly higher expression in TCGA-glioma samples (adjusted *P* < 0.05). Among these, 55 events were located within the genomic regions of “surfaceome” genes with an average expression of TPM ≥ 10 in the TCGA-glioma dataset. However, when defining events with an IRratio > 0.1 and coverage of ≥ 3 reads across the entire intron, as recommended [33], all 55 candidate events were found to be positive in more than 1% of GTEx-brain samples. Moreover, none of these events were present in 10% or more of the cases in any disease groups (IDHwt, IDH-A, or IDH-O) in the TCGA-glioma dataset (**Supplementary Table S3**). As such, due to the absence of tumor-specific IR events, we focused on the exon-exon junction events hereafter.

Among the identified 66 tumor-specific AS events, 42 events were predicted to occur within extracellular segments of the proteins.We then performed an *in silico* translation of the nucleotide sequence of the identified events into an amino-acid sequence. Approximately two-thirds of the events (26, 61.9%) were predicted to generate premature termination codons (PTCs) due to frame-shift changes and were excluded from the subsequent analysis. Only events predicted to cause alterations within the extracellular region of the molecule, while retaining intact transmembrane segments, were regarded as “pass-filter” and retained. After careful manual review, along with the positive control *EGFR*vIII, 13 additional events from 7 genes—*BCAN*, *TREM2*, *NRCAM*, *PTPRZ1*, *NCAM1*, *NCAM2,* and *DLL3* passed all the filters and were identified as the most promising candidates (**Fig. 1b**).

**Fig. 1.**
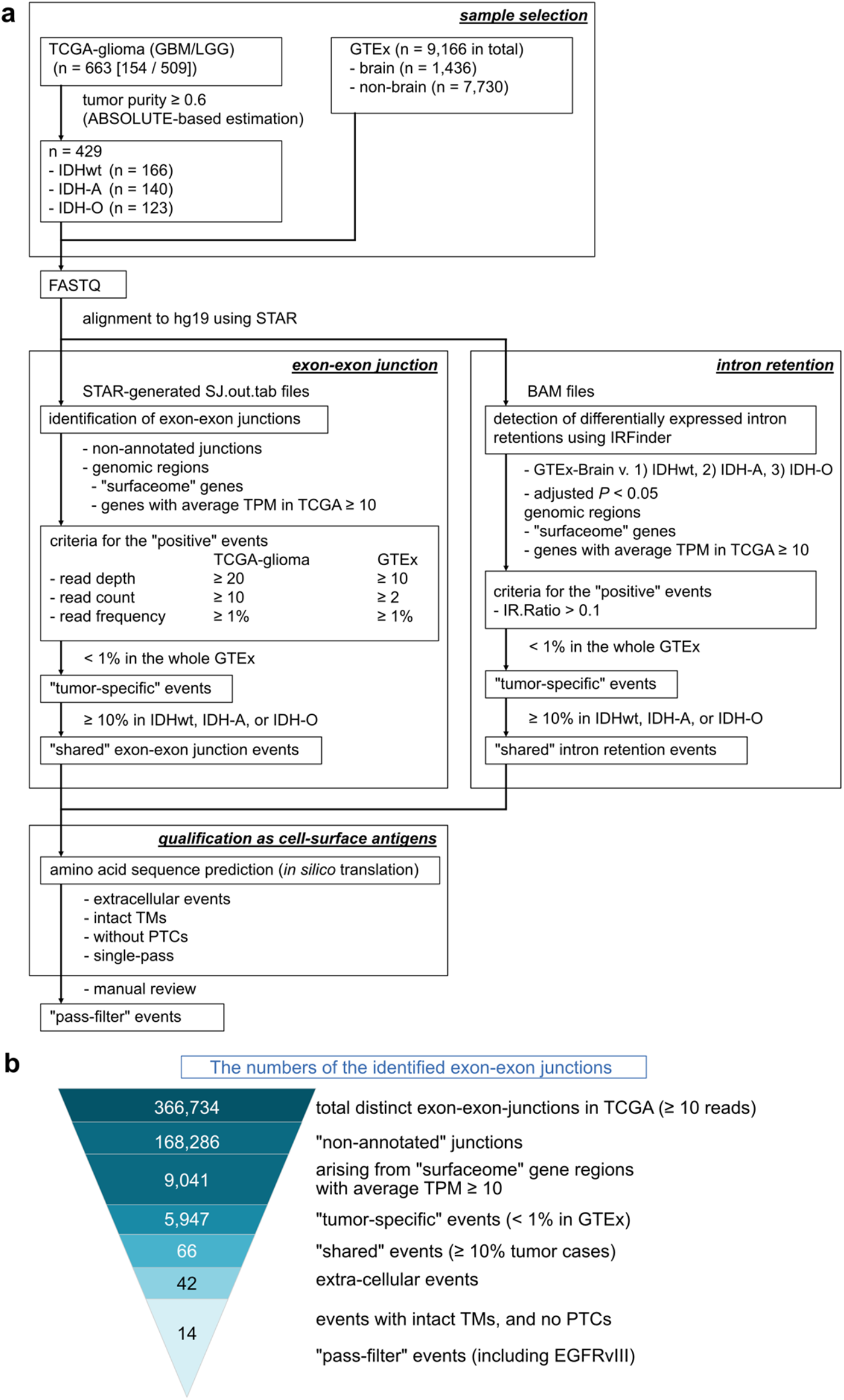
Analysis workflow to identify the tumor-specific AS events as potential sources for cell-surface neoantigens. **a.** Overview of the analytic workflow of The Cancer Genome Atlas (TCGA) glioma and the Genotype-Tissue Expression (GTEx) project datasets. **b.** The number of exon-exon junctions identified in each step of the screening analyses.

### Characteristics of the identified 13 AS events

Among 2,886 surfaceome genes tested, 737 genes had median expression TPM values ≥ 10 in the TCGA-glioma cohort (**Fig. 2a**). Out of the 8 genes above, 6 were identified in the top 60 genes (*BCAN*, average TPM 1,505 [1^st^]; *PTPRZ1*, 589 [13^th^]; *NCAM1*, 249 [29^th^]; *EGFR*, 185 [41^st^]; *NRCAM*, 169 [45^th^]; *DLL3*, 144 [54^th^]). All eight genes were significantly highly expressed in TCGA-glioma samples compared with the GTEx normal tissue samples (**Fig. 2b** **and Supplementary Fig. S1**).

For each of the 13 events and *EGFR*vIII, the identified AS pattern, the abundance of the supportive read count, and the frequency are displayed in **Fig. 2c–p**. The patterns included alternative 3’ splice-site (n = 7 events), alternative 5’ splice-site (n = 4), exon skipping (n = 1, “*NRCAM*-01”), and cryptic intron (n = 1, “*PTPRZ1*-04”). Reflecting the higher gene-level expressions, most read counts were substantially higher in TCGA-glioma samples than in the GTEx normal tissue samples. Certain events, such as *BCAN*-01 and *PTPRZ1*-02, exhibited junction read frequencies of approximately 10% in many “positive” cases. Conversely, the rest of the events displayed a junction-supportive read frequency between 1% and 5% across the majority of the TCGA-glioma cases. A positive control event *EGFR*vIII exhibited an outstandingly higher frequency spanning 50–90%, highlighting its amplification status [32].

Regarding the positive sample rate, each event exhibited distinct patterns among the disease groups. For instance, *BCAN*-01, *BCAN*-03, *NRCAM*-01, *NCAM*-01, *DLL3*-01, -02, and *NCAM2*-01 were more frequently positive in the IDH-O or IDH-A and IDH-O groups. In contrast, *PTPRZ1*-03 was predominantly observed in the IDHwt group, similar to *EGFR*vIII. Furthermore, certain events displayed a relatively high positive sample rate across the disease groups in common. For instance, *BCAN*-01 was positive in 15.7% of IDHwt, 33.6% of IDH-A, and 42.3% of IDH-O cases, with a rate of 0.2% in GTEx samples. Similarly, *PTPRZ1*-02 exhibited positivity in 18.1% of IDHwt, 16.4% of IDH-A, and 24.4% of IDH-O cases, while the rate was 0.15% in GTEx samples. These findings collectively suggest the potential of an off-the-shelf approach in future therapeutic development, although inter-disease differences should also be thoroughly considered.

Moreover, we analyzed additional transcriptome (bulk RNA-seq) data of patient-derived xenograft-origin glioblastoma (PDX-GBM) cell lines [34] both for validation purposes as well as screening purposes for the subsequent experimental investigation. Among the candidates mentioned above, 10 out of 13 were found to be present, with 9 present in multiple cases, along with *EGFR*vIII (**Supplementary Fig. S2**). Overall, the analysis of this external dataset provided further evidence supporting that these candidate AS events are present in other datasets.

**Fig. 2.**
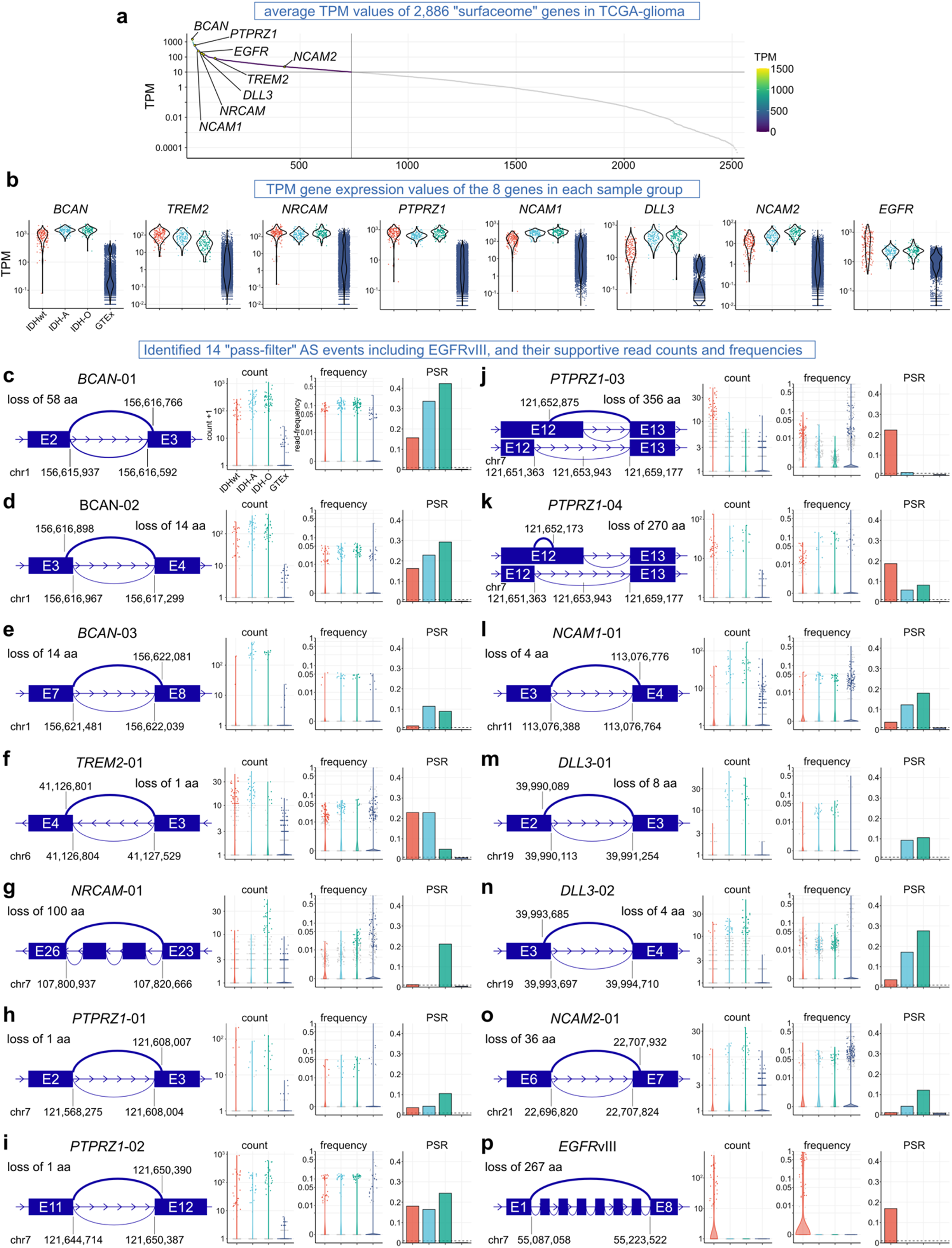
Characteristics of the identified candidate cell-surface tumor-specific AS events. **a.** Dot plots illustrating the gene-level expression of cell-surface (“surfaceome”) genes in the TCGA-glioma dataset. Genes with an average expression of TPM ≥ 10 are depicted in color, while those with an average TPM < 10 are shown in gray. The eight genes identified as the final candidates are labeled for reference. **b.** Dot and violin plots depicting the gene level expression of the eight genes in each sample group. Statistical analysis data is provided in **Supplementary Fig. S1**. **c–p,** Details of the 13 identified events and *EGFR*vIII as a positive control. Each event is accompanied by a schematic illustration of the identified AS pattern, total supportive read count, supportive read frequency, and positive sample rate (PSR) in each sample group. The numbers in the schema indicate the genomic coordinates of the junction edges on the human genome hg19. Note that the Y-axis scales for the ‘count’ panels vary across the events, while those for ‘frequency’ and PSR remain consistent.

### Validation of predicted cryptic transcripts using full-length amplicon sequencing

An inherent shortcoming of ordinal bulk RNA-seq is the inability to determine full-length transcript sequences solely based on short-read sequencing, especially for isoforms with high complexity [35–37]. To address this limitation and provide a foundation for further investigations, we conducted full-length amplicon sequencing using the MinION, a nanopore sequencer from Oxford Nanopore Technologies, on eight PDX-GBM cell lines [34]. We chose to focus on *PTPRZ1* for this experiment because of its robust expression and the highest number of predicted AS events (n = 4) in the candidate list.

We designed the PCR primer pair to bind the 5’-and 3’-untranslated regions (UTRs), allowing us to amplify the entire coding sequence (CDS) (**Fig. 3a**). Following the PCR amplification, we observed various amplicon lengths, in addition to the expected size compatible with canonical transcript *PTPRZ1*-002 (ENST00000449182, 4,875 bp), on agarose gel electrophoresis (**Fig. 3b**). After purification, we prepared the amplicon library, barcoded, pooled, and sequenced the samples using a MINION sequencer. Consistent with the observation on the agarose gel electrophoresis, the subsequent analysis revealed the distribution of the various lengths of the sequencing reads mapped on hg19 chromosome 7, where the *PTPRZ1* gene is located (**Fig. 3c**). We further analyzed the acquired data using the FLAIR platform [38]. This analysis identified 357 different isoforms, each supported by 3 or more sequencing reads. Among these, the top 50 most frequently identified isoforms are displayed in **Fig. 3d** and **Supplementary Fig. S3**.

In all eight samples, the most frequently detected isoform was the canonical isoform *PTPRZ1*-002 with exon 16 (ENSE00001288312) present, termed *PTPRZ1*-002’ (average, 35.6%; range, 25.2–43.5%). The second most frequent was *PTPRZ1*-002 (average, 20%; range, 12.1–29.3%). The other canonical isoform, *PTPRZ1*-001 (ENST00000393386), was found to be the 11^th^ most frequent (average, 1.2%; range, 0.04–6.3%). On the other hand, the two most frequently detected non-canonical isoforms were *PTPRZ1*-002’ containing AS-02 (average, 8.3%; range, 5.8–9.9%) and *PTPRZ1*-002’ containing AS-01 (average, 5.5%; range, 4.0–6.5%). Additionally, as shown in **Fig. 3e–f**, other identified non-canonical isoforms include those harboring both AS-01 and AS-02 (7^th^ most frequent), exon-skipping of the exon 2 (8^th^), IR of the intron 7 (9^th^), and AS-01,-02, and 03 (31^st^). The complete list of the top 50 most frequent isoforms is provided in **Supplementary Fig. S3** with additional details. Notably, all the four predicted AS events (*PTPRZ1*-01, -02, -03, and -04) were detected either individually or in combination with other AS events (average, -01, 14.0%; -02, 19.8%; -03, 0.3%; -04, 1.3%) (**Fig. 3e–f**). Intriguingly, this full-length amplicon sequencing also revealed the co-occurrence of multiple AS events within a single transcript, which is challenging to discern solely through short-read sequencing.

Taken together, our full-length amplicon sequencing assay substantiates the presence of several candidate AS events predicted based on short-read sequencing. Furthermore, the co-occurrence of multiple AS events underscores their complexity and highlights the utility of long-read sequencing-based investigation in identifying and characterizing cryptic isoforms.

**Fig. 3.**
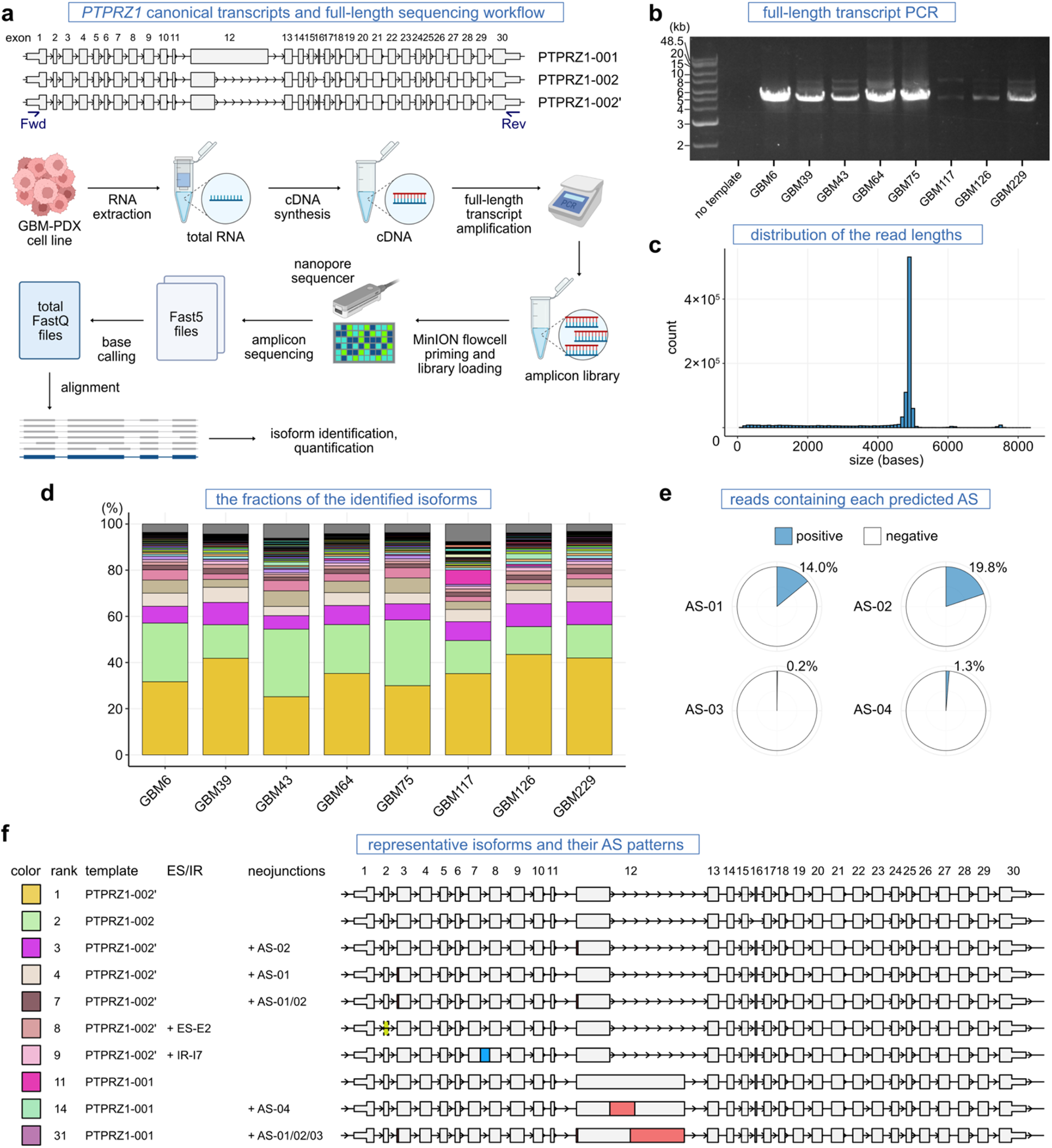
The presence of cryptic transcripts demonstrated through full-length amplicon sequencing. **a.** Canonical transcripts of *PTPRZ1* and the experiment workflow of nanopore full-length amplicon sequencing. In the figure, Fwd and Rev indicate primer-binding sites designed within 5’-and 3’-untranslated regions (UTRs), respectively. **b.** Agarose gel electrophoresis of the *PTPRZ1* full-length transcript amplicon. **c.** Histogram showing the distribution of the read lengths mapped on the hg19 chromosome 7 where the *PTPRZ1* gene is located. **d.** Bar plot depicting the percentage of read counts for each identified transcript. The top 50 most frequently identified isoforms are highlighted in color, while the remaining are shown in gray. The legend for the color code is provided in **Supplementary Fig. S3**. **e.** Pie chart displaying the percentages of total reads of all isoforms involving each candidate AS event. **f.** Schematic illustration of representative canonical and cryptic isoforms found in the top 50 isoforms. The colors in the left square correspond to Fig. 3e. The rank number indicates the ranking in the list. Predicted AS patterns (*PTPRZ1-*01, -02, -03, and -04) (red), and other exon skipping (yellow), intron retention (blue), and cryptic intron (purple) events are highlighted.

### Spatiotemporal heterogeneity of tumor-specific AS events in clinical tumor samples

Spatial, temporal, inter-patient, and intra-patient heterogeneity are among the most prominent characteristics of glioma, which makes the disease more difficult to treat [18,39–42]. However, in contrast to the well-studied mutational landscape, our understanding of the AS landscape remains limited. This knowledge gap is particularly critical in antigen-oriented immunotherapy, where high heterogeneity increases the risk of antigen loss and treatment failure. To this end, we explored two unique datasets of clinical sample transcriptomes: 1) those collected through spatially mapped multi-sampling and 2) those obtained longitudinally from primary and recurrent paired tumor samples within the same individual patients.

In the first dataset, we examined transcriptomes of 169 tumor pieces from 24 primary gliomas (3– 12 pieces with sufficient tumor purity per patient, averaging 7 pieces) (**Fig. 4a–b, Supplementary Table S4**). Notably, intratumoral heterogeneity was prominent in most cases and events. Tumor-wide (100%) positivity was exhibited only for four event-case pairs (*PTPRZ1*-03 in tumor T2; *NCAM1*-01 in T24; *DLL3*-02 in tumors T15 and T16). While certain events, such as *BCAN*-02, *PTPRZ1*-03, and *DLL3*-02, displayed relatively homogenous positivity, many others were highly heterogeneous. Interestingly, even *EGFR*vIII, one of the most common genetic rearrangement mutations in IDHwt gliomas, exhibited non-negligible intratumoral heterogeneity—three tumors (T1, T2, T3) displayed positivity in multiple tumor pieces, yet none of them were tumor-wide.

In the second dataset, we investigated the transcriptomes of 23 tumors, encompassing 11 pairs/sets of primary and recurrent tumors, including 4 with multisampling tissue collections (**Fig. 4c–d, Supplementary Table S5**). As shown in **Fig. 4d**, when defining the positivity as ≥ 50%, only 10 events (event-patient pairs) were shared between primary and recurrent tumors in this analysis, and most of the detected events were transient—either disappeared or newly emerged at the time of recurrence.

Our investigations of clinical samples reveal remarkable spatiotemporal heterogeneity in candidate AS events, underscoring the challenges associated with the development of AS-derived antigen-targeting therapeutics.

**Fig. 4.**
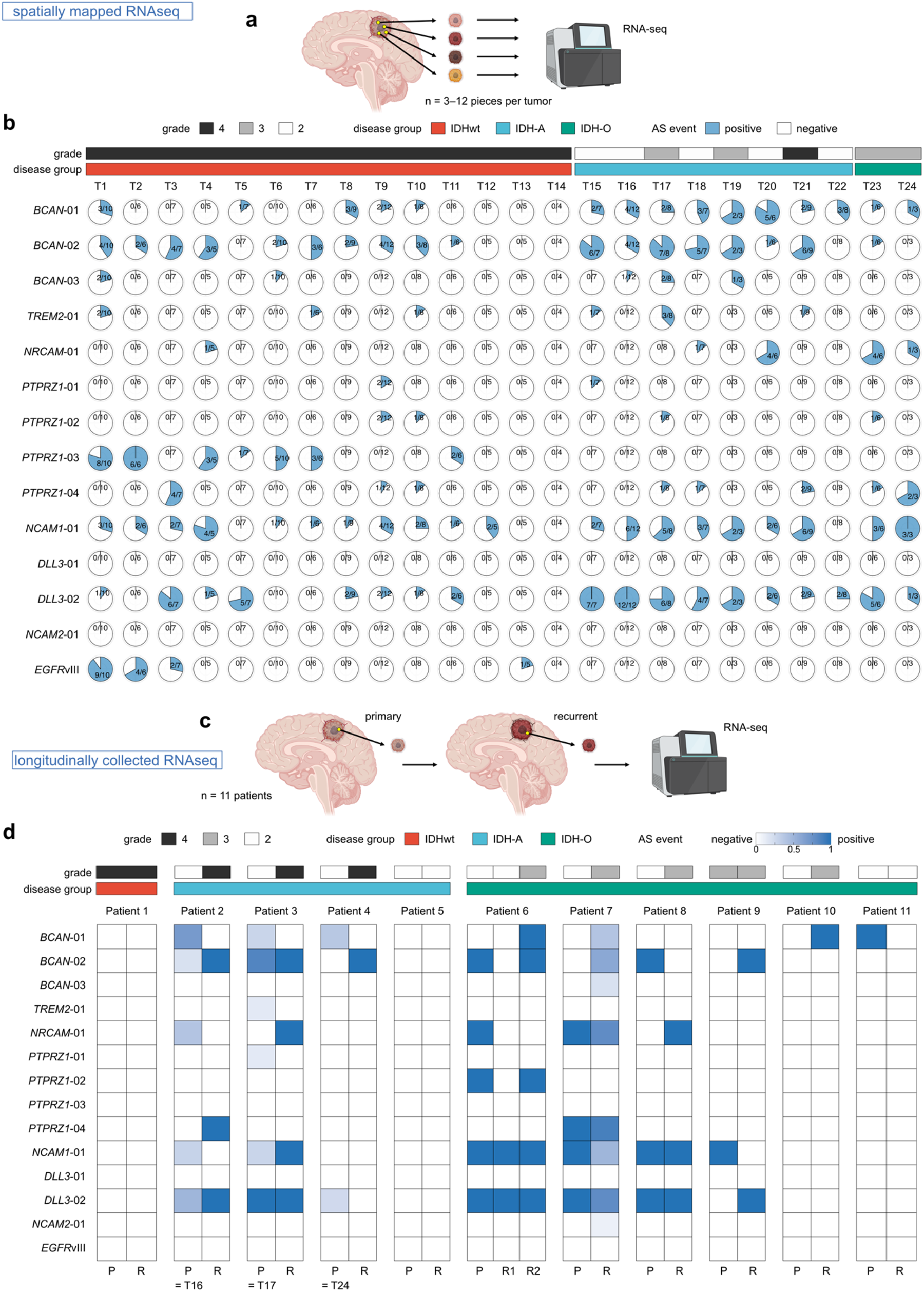
Spatiotemporal heterogeneity of tumor-specific AS events in clinical tumor samples. **a.** Overview of spatially-mapped tumor sample data collection. **b.** Chart displaying the detection of candidate cell-surface AS events in the dataset. The pie charts illustrate the frequency of event detection within the same tumor. Diagnostic information is indicated in the top bars. **c.** Overview of primary and recurrent paired tumor sample data collection. **d.** Heatmap illustrating temporal changes and commonalities of each candidate event in primary and recurrent paired tumor samples.

### CPTAC proteomics data analyses fail to detect candidate AS-derived peptide signals

Protein-level expression is pivotal for antigens to elicit host immunity and to be targeted for immunotherapy. However, transcripts harboring alterations are more likely to undergo degradation before translation, often through nonsense-mediated mRNA decay [43]. Therefore, validating their protein-level expression is crucial. To address this question, we re-analyzed a publicly available proteome dataset from the Clinical Proteomic Tumor Analysis Consortium (CPTAC) [44].

Consistent with the original analysis pipeline, we employed the software MSGF+ with a modified human proteome reference file that incorporated all the predicted cryptic protein FASTA sequences. As shown in **Fig. 5a–b**, the analysis successfully detected two distinct peptide spectra matching *EGFR*vIII-specific peptide fragments with statistical significance. In this analysis, however, none of the predicted AS-derived candidates were reliably supported by mass spectral signals. Among those examined, *BCAN*-03 exhibited the lowest E-value and Q-value, but did not reach statistical significance.

We also applied quantitative analysis using the software MASIC [45] to investigate the relative intensity of each detected peptide spectrum. One of the two *EGFR*vIII-specific peptide fragments (KGNYVVTDHGSCVR) displayed a significantly higher average value in tumor samples than normal controls (average: 2.99 v. 1.06, *P* = 0.02) (**Fig. 5c**). Similarly, the other *EGFR*vIII-specific peptide (GNYVVTDHGSCVR) exhibited a similar trend but did not reach statistical significance (average: 2.42 v. 1.59, *P* = 0.12). In contrast, no difference was observed between tumor and normal samples concerning the *BCAN*-03-related peptide candidate (average: 1.43 v. 1.27, *P* = 0.61). Since the wild-type proteins corresponding to the candidate AS events were expressed and detectable within the dataset (**Supplementary Fig. S4**), it seems unlikely that the failure to detect the predicted aberrant peptide signals can be attributed to the lack of absolute wild-type protein expressions. Therefore, the protein-level expression for the predicted AS candidate events was not supported by the proteomics analysis, except for the positive control *EGFR*vIII, despite such a large-scale cohort dataset (n = 99 tumors). These observations underscore the challenges and uncertainties in the discovery of tumor-specific AS-derived cell-surface neoantigens.

**Fig. 5.**
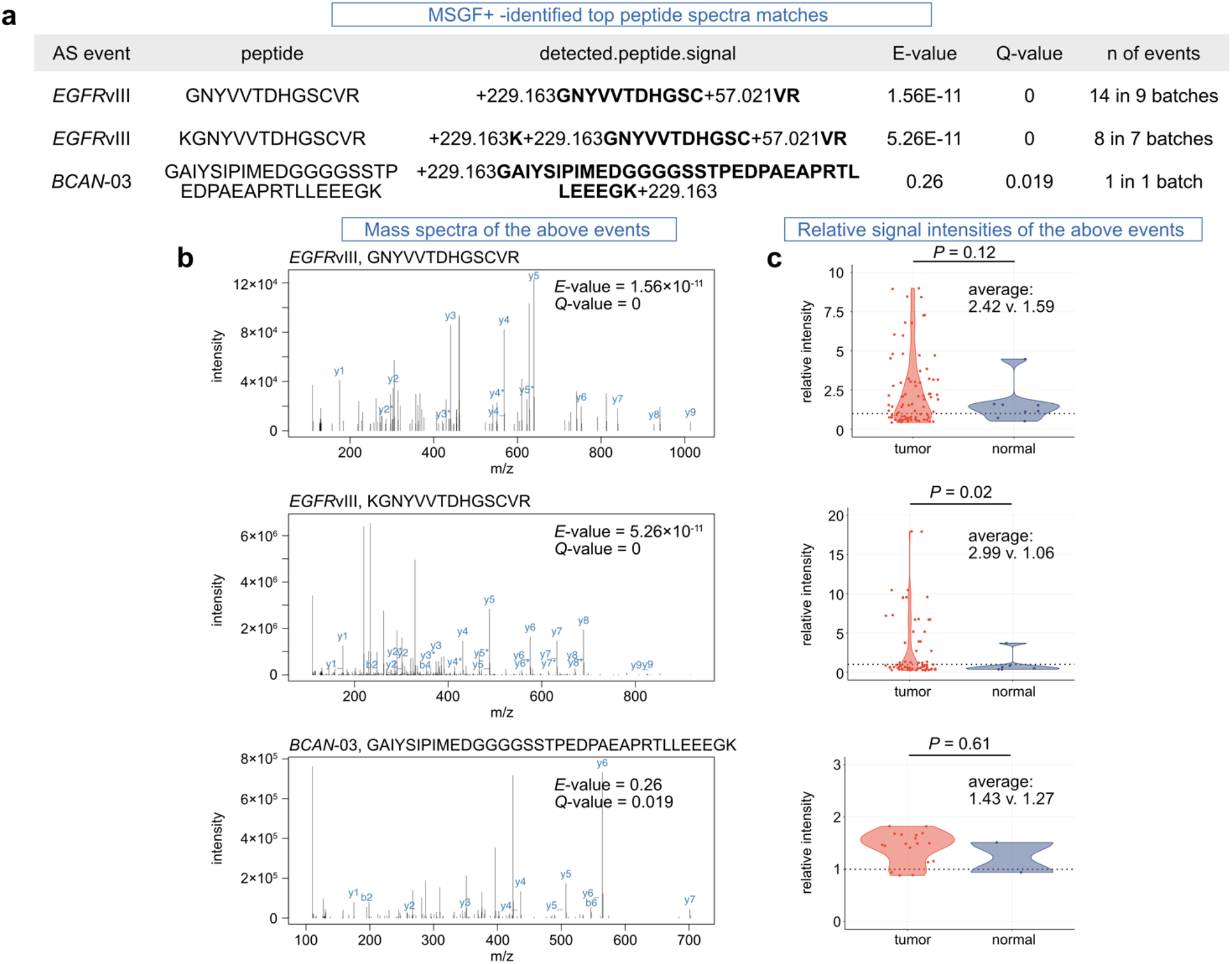
CPTAC proteomics data analyses detect *EGFR*vIII peptide signals but no other candidate AS-derived peptide signals. **a.** MSGF+ result table summarizing the detection of the candidate AS events of our interest. Only top three peptides with the smallest Q-values are shown. **b.** Peptide spectra of two *EGFR*vIII and one *BCAN*-03-derived peptides identified through the analysis with MSGF+. **c.** Dot and violin plots displaying signal intensities of the peptide spectra calculated using MASIC and normalized as relative to the internal reference sample in the dataset. The data of tumor and normal samples are shown separately. *P* values are calculated using *t-*test.

## Discussion

In this study, we embarked on a multi-layered exploration of tumor-specific AS events in glioma. Our primary goal was to uncover a potential reservoir of untapped neoantigens, specifically identifying cell-surface antigen candidates derived from these tumor-specific AS events. The screening analysis comparing the TCGA-glioma and GTEx normal tissue datasets identified 13 tumor-specific AS events as top candidates for further scrutiny. To reinforce our findings, we conducted full-length transcript amplicon sequencing, providing additional evidence of the predicted AS patterns within the expressed transcripts.

Notably, our analysis of unique clinical tumor sample datasets, both spatially mapped and longitudinally collected, revealed a complex landscape characterized by profound spatio-temporal and inter- and intra-patient heterogeneity in the predicted AS events. This heterogeneity, observed across different patients, diverse tumor fragments, and distinct stages of disease progression, emerged as a prominent feature of our investigation. Moreover, through the re-analysis of the CPTAC proteome data, we were unable to corroborate the protein-level expression of the predicted tumor-specific AS-derived neoantigen candidates.

While our pursuit for neoantigen discovery has not yielded the expected results, the findings from our study still hold several valuable implications. First, the analysis of two unique datasets of clinical tumor samples, meticulously curated within our institution, shed light on the intricate transcriptomic diversity of these tumors. The analysis of the spatially mapped tumor dataset revealed the rarity of “tumor-wide” events, with most detected events exhibiting intratumoral heterogeneity, including *EGFR*vIII. Second, temporal heterogeneity was revealed as prominent, with only a limited number of events shared between primary and recurrent tumors within our analyzed dataset. While certain aspects of this heterogeneity may stem from technical constraints intrinsic to sequencing and subsequent analysis, it underscores the significance of recognizing intra- and inter-tumoral heterogeneity in future endeavors of therapeutic development. Otherwise, targeting suboptimal antigens will likely cause clonal evolution and antigen loss [15,19,46], leading to treatment failure. Therefore, irrespective of genomic, transcriptomic, and post-translational events, future antigen discovery efforts should carefully integrate considerations for this spatio-temporal heterogeneity issue, particularly in tumors recognized for their heterogeneity, as exemplified by glioma [18].

Third, a fundamental principle in antigen exploration emphasizes the significance of evaluating protein-level expression. A limitation of this study is that the CPTAC-GBM proteome dataset primarily consists of only glioblastoma cases, with all but one case being IDHwt [44]. Indeed, our transcriptome-based screening analyses indicated a more frequent presence of certain candidate AS events in IDH-mutant tumors. Nevertheless, the absence of peptide signals from these candidate AS-derived peptides, in contrast to the positive control *EGFR*vIII, implies their elusive nature within detectable ranges. In most scenarios, antibody or CAR-T cell recognition positively correlates with the abundance of surface antigens [47]. Although it is crucial to bear in mind that the absence of evidence does not necessarily indicate evidence of absence, our analytical findings did not provide compelling rationale for further pursuit of this avenue in antigen discovery.

Our study additionally highlights the value of long-read sequencing-based investigation using a nanopore platform. Full-length transcript amplicon sequencing revealed a complex landscape of isoforms involving the co-occurrence of multiple AS events, which cannot be accurately determined only through short-read-based sequencing. When interpreting the data, it is imperative to be cautious regarding any quantitative analysis due to the PCR amplification bias introduced as part of the library preparation workflow [48]. Nevertheless, determining the full-length transcript sequence is pivotal for precise antigen prediction since it can directly influence the reading frame, swiveling between an in-frame and frameshift configuration. While our full-length transcript amplicon sequencing focused on *PTPRZ1* as the only target in this study, leveraging a more high-throughput approach could be considered for future antigen discovery attempts involving AS.

Lastly, in the context of antigen discovery exploration or prediction of this nature, it is crucial to always bear in mind that there are no definitive thresholds for ensuring efficacy and safety [7–11,49]. We set arbitrary thresholds, such as a junction-supportive read frequency of less than 1% and a positive sample rate of less than 1% in the GTEx samples, referring to a previous study by Kahles et al. [20]. While we aimed to establish stringent criteria emphasizing tumor-specificity and safety, it is anticipated that analyses with different criteria will identify dissimilar sets of candidates, considering the inevitable trade-off relationship between sensitivity and specificity. Furthermore, despite this stringency, the designation of “less than 1%” does not guarantee an adequate level of safety. Following the discovery phase, the safety profile must be ensured through rigorous assessments of novel antigen candidates.

### Conclusions

In conclusion, our investigation shed light on the challenges in exploring tumor-specific AS-derived cell-surface neoantigens. Their remarkable spatiotemporal heterogeneity and elusive nature at the protein levels underscore a formidable challenge. We should incorporate the lessons learned during this quest as we chart our course. Redirecting our efforts toward intracellular, MHC-presented antigens arising from AS [27] and other tumor-specific post-translational events could offer a more viable avenue. This path may offer more attainable prospects within the expansive landscape of cancer immunotherapy.

## Methods

### Data downloading from the TCGA and GTEx projects

Regarding the TCGA-glioma dataset, raw RNA-seq BAM files of the primary tumor cases found in the study projects TCGA-GBM and TCGA-LGG were downloaded through GDC Data Portal website (https://portal.gdc.cancer.gov/) in February 2019. These BAM files were converted to FASTQ format using Picard (v2.17.1) *SamToFastq* function with default settings. For the GTEx dataset, paired-end FASTQ files were downloaded using pfastq-dump (v0.1.6, github.com/inutano/pfastq-dump) with the ‘*--split-files*’ option. Access permission to protected data on dbGaP was obtained under project number 14627. Clinical, pathological, molecular diagnosis information, and ABSOLUTE-estimated tumor purity data were gathered for the TCGA dataset from the reference [30]. After excluding the samples with unknown or low tumor purity (ABSOLUTE-estimated tumor purity < 0.6) and those lacking essential molecular diagnosis information, 429 tumor samples were retained for this study (115 and 314 tumor cases from TCGA-GBM and TCGA-LGG cohorts, respectively). Based on the *IDH1/2* mutation and chromosome 1p/19q statuses, the tumor samples were classified into glioma IDH wildtype (IDHwt), astrocytoma, IDH-mutant, 1p/19q-non-codeleted (IDH-A), or oligodendroglioma, IDH-mutant, 1p/19q-codeleted (IDH-O) [31]. For the GTEx dataset, annotation information for organs and tissue types was obtained from GTEx_v7_Annotations_SampleAttributesDS.txt, available on the GTEx Data Portal website (https://www.gtexportal.org/home/downloads/adult-gtex#metadata). RNA-seq gene expression abundance data of both the TCGA and the GTEx datasets were downloaded through the UCSC Xena-Toil web portal (dataset ID: TcgaTargetGtex_rsem_gene_tpm; filename: ‘TcgaTargetGtex_rsem_gene_tpm.gz’; version: 2016-09-03) [50]. The downloaded log2(transcript-per-million [TPM] + 0.001) values were transformed back to original TPM values for subsequent quantitative analyses and visualization.

### RNA sequencing read alignment

A genome index was built using STAR (v2.7.7a) and a reference genome ‘GRCh37.primary_assembly.genome.fa’ downloaded from GENCODE, along with ‘Homo_sapiens.GRCh37.87.chr.gtf’ from Ensembl. The following command was used: *STAR -- runMode genomeGenerate --genomeDir $GENOME –genomeFastaFiles $FASTA --sjdbGTFfile $GTF - -sjdbOverhang 100 --runThreadN $N*. Then, FASTQ reads were aligned to the human reference genome hg19, generating BAM files and SJ.out.tab files for each sample. The alignment process followed the command reported by Kahles et al.[20]: *STAR --genomeDir $GENOME --readFilesIn $FQ1 $FQ2 -- runThreadN $N --outFilterMultimapScoreRange 1 --outFilterMultimapNmax 20 -- outFilterMismatchNmax 10 --alignIntronMax 500000 --alignMatesGapMax 1000000 --sjdbScore 2 -- alignSJDBoverhangMin 1 --genomeLoad NoSharedMemory --limitBAMsortRAM 70000000000 -- readFilesCommand cat --outFilterMatchNminOverLread 0.33 --outFilterScoreMinOverLread 0.33 -- sjdbOverhang 100 --outSAMstrandField intronMotif --outSAMattributes NH HI NM MD AS XS -- limitSjdbInsertNsj 2000000 --outSAMunmapped None --outSAMtype BAM SortedByCoordinate -- outSAMheaderHD @HD VN1.4 --twopassMode Basic --outSAMmultNmax 1*. Samtools (v.1.10) was used for sorting and indexing the generated BAM files. The resulting BAM files and the SJ.out.tab files containing detected junction information were utilized for subsequent analyses.

### Identification of “surfaceome” genes

To investigate the genes encoding cell-surface proteins, we selected 2,886 “surfaceome” genes, referring to [51]. Amino acid position information of the extracellular segment was also obtained from the same database. The genomic coordinates of each gene were obtained from the GTF files described above, and were referred to for matching of the junction coordinates.

### Identification of the exon-exon splicing junctions

Quantitative information regarding the high-confidence collapsed splice junctions was extracted from the STAR-generated SJ.out.tab files. A unique junction ID was defined based on the information in columns 1–4 (chromosome, first base of the intron, last base of the intron, and the strand). The column 7 provided the count of uniquely mapped reads crossing the junction, allowing for the acquisition of quantitative data for each unique splicing junction.

The positivity of a junction in each sample was determined based on the following three factors: 1) the junction supportive read count; 2) the read depth at the junction; and 3) the junction supportive read frequency. The junction supportive read count represents the number of uniquely mapped reads as described earlier. For each investigated junction, the most dominant overlapping junction was defined either by the most abundant ‘annotated’ junction or, if no ‘annotated’ junctions were present, by the most abundant junction in the total TCGA-glioma samples. For each junction, the selected same most dominant overlapping junction was applied to both the TCGA-glioma and the GTEx datasets. The read depth was calculated as the sum of the junction read count and the corresponding most dominant overlapping junction. The junction supportive read frequency was obtained by dividing the junction-supportive read counts by the read depth. Due to substantial differences in RNA-seq data size and detected total junction read counts between the TCGA-glioma and the GTEx datasets (the total read count in RNAseq data: 1.3×10^8^ and 8.8×10^7^, respectively; t-test *P* = 5.5×10^-118^; the total number of uniquely mapped junction reads on average: 6.1×10^6^ and 4.7×10^6^, respectively; t-test *P* = 7.5×10^-5^; **Supplementary Fig. S5a–b**), different cut-off values were applied to determine the positivity for each event. For the TCGA-glioma dataset, junctions with 1) junction supportive read counts ≥ 10, 2) read depth at the junction ≥ 20, and 3) junction-supportive read frequency ≥ 0.01 (1%) were considered positive for the event. In the GTEx dataset, junctions with 1) read counts ≥ 2, 2) read depth ≥ 10, and 3) read frequency ≥ 0.01 were deemed positive for the event. For each identified distinct event, the positive sample rate (PSR) was determined by dividing the number of positive samples by the total number of samples in the group (n = 166 for IDHwt; n = 140 for IDH-A; n = 123 for IDH-O; n = 9,166 for all GTEx samples). Events with PSR < 0.01 (1%) in the GTEx samples were defined as tumor-specific, while those with PSR ≥ 0.1 (10%) in at least one disease group (either IDHwt, IDH-A, or IDH-O) were considered shared events.

### Intron retention analysis

For the detection of intron retention (IR)-type AS events, the RNA-seq BAM files of both the TCGA-glioma (n = 429) and the brain samples of the GTEx datasets (n = 1,436) were analyzed using IRFinder (v1.2.3) [33]. The command used to generate the output file ‘IRFinder-IR-nondir.txt’ was as follows: *IRFinder -m BAM -r REF/Human-hg19-release75 -d $DIRECTORY $BAM*.

Differential IR analysis was conducted by passing the above IRFinder result files to DESeq2 [52] in the R interface. The fold change of IR between the two groups were tested using a generalized linear model (GLM) based on the provided design matrix, following the software recommendation [33]. The analyses were repeated by comparing the 1,436 brain samples in the GTEx dataset and 1) IDHwt (n = 166); 2) IDH-A (n = 140); 3) IDH-O (n = 123). In each comparison, events that were more frequent in tumor samples (positive fold change values) with adjusted *P* values < 0.05 were considered significantly differentially expressed and retained. In the subsequent investigation, events with an IRratio > 0.1 and at least coverage of three reads across the entire intron were defined as positive in the sample, as recommended by the developer [33].

### In silico translation into amino acid sequences

For each candidate AS event, one or more canonical isoforms were identified as the templates. For instance, in the case of *PTPRZ1*, which has two canonical isoforms (ENST00000393386 [8,175 bp] and ENST00000449182 [4,875 bp]), both were used as templates for *PTPRZ1*-related candidate AS events. The altered nucleotide sequence for each candidate AS event was predicted based on the genomic coordinates of the event and the nucleotide sequence of the corresponding template transcripts. This process utilized the R packages ‘GenomicRanges,’ ‘ensembldb,’ ‘EnsDb.Hsapiens.v75,’ and ‘BSgenome.Hsapiens.UCSC.hg19’. The predicted nucleotide sequences with alterations were translated into amino acid sequences using the *DNAString()* and *translate()* functions in the R package ‘Biostrings’. Any termination codons, including the PTCs, were marked by asterisks (*) in the output. The same *in silico* translation was also performed with the wildtype sequence and cross-validated with the sequence information in the Ensembl database to ensure accuracy.

### Filtering steps to identify non-annotated, highly expressed, cell-surface, tumor-specific exon-exon junctions

The following filtering steps were employed to identify the non-annotated, tumor-specific, expressed, and surfaceome-gene-derived AS events:

1. Examination of all exon-exon splicing junctions with a junction-supportive read count of ≥ 10 in the 429 samples of the TCGA-glioma cohort, resulting in the listing of all unique junction IDs (n = 366,734 unique junctions).
2. Exclusion of junctions found in the reference junction database generated by STAR based on the reference GTF file, retaining only the “non-annotated” junctions (n = 168,286).
3. Retention of junctions overlapping with any portion of the genomic coordinates of the surfaceome genes that have an average gene expression TPM value ≥ 10 in the TCGA-glioma dataset (n = 9,041).
4. Retention of events with a PSR < 0.01 (1%) in the GTEx samples, labeled as “tumor-specific” (n = 5,947).
5. Retention of events with a PSR ≥ 0.1 (10%) in either IDHwt, IDH-A, or IDH-O, categorized as “shared” events (n = 66).
6. Retention of events affecting the extracellular segments, based on the extracellular segment information from the surfaceome database (n = 42).
7. Evaluation of the predicted altered amino acid FASTA sequence information for each pass-filter event, considering whether the consequence is in-frame or frameshift, whether PTCs are induced, and whether the transmembrane segment is intact. Because of the complexity of the prediction, multi-pass proteins (those with multiple transmembrane segments) were excluded. Events predicted to be without PTCs and with intact transmembrane segments were retained (n = 14).
8. Manual examination and confirmation of these pass-filter events.

### Validation analysis with an external dataset

For both in silico and experimental validations, we obtained raw bulk RNA-seq data of glioma patient-derived xenograft (PDX) models (n = 66 tumors) from the Mayo Clinic GBM PDX National Resource [34] via the NCBI Gene Expression Omnibus (GEO) under Accession PRJNA548556. Paired-end FASTQ files were downloaded using pfastq-dump (v0.1.6) with the following option ‘—disable-multithreading --split-3 *--split-files -- s $SRA_ID*’. The processing of the downloaded FASTQ files, quantification of the splicing junctions, and assessment of the presence of the junctions in each sample were all performed in the same manner as for the TCGA-glioma dataset.

### Tumor cell line culture

From the Mayo Clinic GBM PDX National Resource [34], we obtained the following 8 GBM-PDX cell lines under the material transfer agreement: GBM6, 39, 43, 64, 75, 117, 126, and 229. These cells were maintained in complete NeuroCult media, which consisted of NS-A Basal Medium (Human) (STEMCELL Technologies, 05750) supplemented with 10% NeuroCult™ Proliferation Supplement (Human) (STEMCELL Technologies, 05753), heparin solution (final concentration, 2 µg/mL, STEMCELL Technologies, 07980), recombinant human EGF (20 ng/mL, PeproTech, AF-100-15), recombinant human FGF-basic (20 ng/mL, PeproTech, 100-18B), and 1% Penicillin-Streptomycin (Gibco, 15070063). The cells were cultured in spheroid form on culture dishes with an ultra-low attachment surface (Corning, 3471, 3473, 3814, or 4616) and maintained in a humidified incubator at 37°C with 5% CO2. Passaging of the cells was performed by dissociating them using Accutase (Innovative Cell Technologies, AT104) and then plating them onto new culture dishes. EGF and FGF-basic were directly supplemented to the media twice a week during maintenance. Regular testing for mycoplasma infection was conducted every 3–4 months using the PlasmoTest mycoplasma detection kit (InvivoGen, catalog # rep-pt1). No other authentication assay was performed.

### RNA extraction and cDNA synthesis

Total RNA was extracted from tumor cell culture pellets using the RNeasy Mini Kit (Qiagen, 74106) and the RNase-Free DNase Set (Qiagen, 79254) following the manufacturer’s protocol. The RNA integrity was evaluated using the Agilent Bioanalyzer 2100 and confirmed to be RIN ≥ 7 (range, 7.3–9.9). Subsequently, complementary DNA (cDNA) was synthesized from the total RNA samples using the TaKaRa PrimeScript 1st strand cDNA Synthesis Kit (TaKaRa, 6110A).

### Acquisition of *PTPRZ1* full length transcript amplicons

To obtain the amplicons of the entire coding sequence (CDS) region of *PTPRZ1* transcripts, the following oligo primers were designed to bind to the 5’-UTR and 3’-UTR of *PTPRZ1* transcripts, respectively: Fwd: 5’-ACCGTCTGGAAATGCGAATC-3’; Rev: 5’-TGGTCCATGGAGACACTAGG-3’. Polymerase chain reaction (PCR) was performed using PrimeSTAR GXL DNA Polymerase (TaKaRa, R050A), following the recommended Rapid PCR protocol, consisting of 35 cycles of 98℃ for 10 sec, 60℃ for 15 sec, and 68℃ for 80sec. The amplicons (PCR products) were imaged after electrophoresis on a 0.8% agarose gel. Gel pieces of each lane were excised from the agarose gel, digested, and purified using the NucleoSpin® Gel and PCR Clean-Up kit (TaKaRa, 740609), according to the manufacturer’s protocols.

### Nanopore full-length transcript amplicon sequencing

The library preparation of the nanopore full-length transcript amplicon was conducted following the protocol “native-barcoding-amplicons-NBA_9093_v109_revD_12Nov2019-minion,” provided by Oxford Nanopore Technologies (ONT). The following materials were used: Ligation Sequencing Kit (ONT, SQK-LSK109); Native Barcoding Expansion 1-12 (PCR-free) (ONT, EXP-NBD104); Flow Cell Priming Kit (ONT, EXP-FLP002); Agencourt AMPure XP beads (Beckman Coulter, A63880); Blunt/TA Ligase Master Mix (New England Biolabs, M0367S); NEBNext Quick Ligation Reaction Buffer (New England Biolabs, B6058S); NEBNext Companion Module for Oxford Nanopore Technologies Ligation (New England Biolabs, E7180S). As per the recommended protocol, the sample concentration was measured using the Qubit™ dsDNA HS Assay Kit (Invitrogen, Q32854) and Qubit Fluorometer (Invitrogen), and was adjusted accordingly at each step. After barcode ligation, all the individual libraries were pooled. Subsequently, following adapter ligation and the concentration measurement, 50 fmol of the pooled library was loaded to the R9.4.1 flow cells (ONT, FLO-MIN106D) on the MinION Mk1B sequencer.

### Analysis of the full-length transcript amplicon sequencing

The sequencing data was acquired using the MinKNOW software (v4.0.20) with high-accuracy mode. The generated Fast5 data were analyzed, excluding the Fast5_fail data and only focusing solely on the Fast5_pass data. Base calling was performed using Guppy basecaller (4.2.2+effbaf8) with the following command: *guppy_basecaller --flowcell FLO-MIN106 --kit SQK-LSK109 --cpu_threads_per_caller $NUM_CPU_THREADS --num_callers $NUM_CPUS --compress_fastq --barcode_kits EXP-NBD104 -- trim_barcodes --qscore_filtering -i $INPUT_DIR -s $OUTPUT_DIR*.

The generated FASTQ files were analyzed using FLAIR (v2.0.0) [38] in the Python 3.9.16 environment. For the FLAIR analyses, a customized reference genome was prepared by extracting the chromosome 7 region where *PTPRZ1* gene is located in the human genome hg19 (GRCh37). Additionally, the GTF annotation file was also narrowed down to the chromosome 7 region from the file ‘Homo_sapiens.GRCh37.87.chr.gtf,’ and each of the expected AS patterns of *PTPRZ1* was concatenated.

The first module, ‘flair align,’ was performed for the FASTQ files with the reference genome, generating BED files. The second module, ‘flair correct,’ was performed with the BED files using the following command: *flair correct -q $BED -g $GENOME -t $N -f $GTF -o $OUT_PREFIX*. The third module, ‘flair collapse,’ was conducted with the concatenated single FASTQ file and the concatenated single BED file of all tested samples using the following command: *flair collapse -g $GENOME -q $BED -r $FQ -t $N -f $GTF -o $OUT_PREFIX --keep_intermediate --generate_map --check_splice -- annotation_reliant generate --trust_ends*. Based on the defined high-confidence isoforms, the fourth module, ‘flair quantify,’ was performed to obtain the read count data of each isoform in each sample using the following command: *flair quantify -r $MANIFEST -i $ISOFORM -o $OUT_PREFIX2 -t $N -- generate_map --sample_id_only --trust_ends*. The output data were summarized and visualized in the R 4.1.3 environment.

### Clinical sample data acquisition and analysis

Detailed information on the acquisition and processing of clinical tumor sample has been previously documented [53]. For the dataset of spatially mapped tumor samples, tumors with a T2-defined lesion of 4-5 cm diameter or greater, enabling sufficient capacity for multi-sample collection, were considered eligible for inclusion. During tumor resection craniotomy, spatially mapped samples were collected by neurosurgeons to represent a comprehensive, randomly sampled view of the total tumor geography. The 3D coordinates of each resected tumor piece were recorded and registered to a preoperative magnetic resonance image (MRI) using a Brainlab Cranial Navigation system (BrainLAB, v3). The weights of the collected samples ranged between 75–225 mg. When feasible, 2/3 of each sample were flash-frozen in liquid nitrogen for DNA and RNA extraction and subsequent sequencing, while the remaining 1/3 was utilized for histopathological examinations.

Genomic DNA and RNA were simultaneously extracted from frozen tissues using an AllPrep DNA/RNA/miRNA Universal kit (Qiagen, 80224), following the manufacturer’s instructions. Whole-exome sequencing (WES) was conducted as previously reported [39]. Tumor-purity was estimated for each sample using either FACETS [54] for IDH-wt and IDH-O cases or the variant allele frequency (VAF) of *IDH1* R132 mutations for IDH-A cases, based on WES data. RNA-seq libraries were prepared using the KAPA Stranded mRNA-Seq kit (Kapa Biosystems, KK8421) or KAPA mRNA HyperPrep Kit (Kapa Biosystems, KK8580), a streamlined version of Stranded mRNA-Seq kit, and sequenced on a HiSeq2500, 4000, or NovaSeq (Illumina). The sequencing FASTQ files were processed and analyzed in the same manner as the TCGA and GTEx datasets, as described earlier. Samples with estimated tumor purity ≥ 50% in IDHwt and IDH-O cases, and the VAF of *IDH1* R132 > 25% in IDH-A cases, were included in the analysis of the spatially mapped dataset. For the longitudinally collected tumor dataset, tissue collections, processing, and analyses were carried out as previously reported [39].

The RNA-seq data of the spatially mapped dataset have been deposited in the European Genome-phenome Archive (EGA) under accession number EGAS00001003710. The data for the longitudinally collected dataset have been deposited in the EGA under accession number EGAS00001002368 and dbGaP study accession: phs002034.v1.p1.

### Data downloading of the CPTAC-GBM dataset

The CPTAC-GBM [44] human glioblastoma proteome raw files (.raw), the processed mass spectra data files (.mzML), and the metadata files were obtained in April 2020 through the NIH Proteomic Data Commons (Study ID PDC000204). The download included 264 separate files from 11 run batches. Additionally, quantitative analysis data, specifically the reporter ion intensity Log2Ratio of the identified proteins, was also downloaded (filename: cfe9f4a2-1797-11ea-9bfa-0a42f3c845fe-log2_ratio).

### Peptide spectra database search using MSGF+

For the proteomics database search, the reference human proteome database (filename: UP000005640_9606.fasta) was downloaded from UniProt in November 2020. The FASTA sequence of the altered proteins putatively derived from the 14 candidate AS events, including *EGFR*vIII, was concatenated and used as the customized reference proteome database. Following the original workflow of the CPTAC-GBM project, peptide spectra database search was performed using MSGF+ (v2020.07.02) [55] in a Java environment (v1.8.0_382) with the following configurations: *PrecursorMassTolerance=20ppm; IsotopeError: 0, 1; NumMods=3; FragmentationMethodID=0; InstrumentID=1; EnzymeID=1; NTT=2; TDA=1; NumThreads=All; MinPepLength=6; MaxPepLength=50; MinCharge=2; MaxCharge=5; NumMatchesPerSpec=1; MaxMissedCleavages=1; StaticMod=229.1629, *, fix, N-term, TMT11plex; StaticMod=229.1629, K, fix, any, TMT11plex; StaticMod=C2H3N1O1, C, fix, any, Carbamidomethyl; DynamicMod=O1, MP, opt, any, Oxidation*. The database search was performed with the following command: *java -Xmx3500M -jar MSGFPlus.jar - s $MZML -d $FASTA -conf $CONFIG -o $MZID*.

### Quantification analysis using MASIC

From the proteome raw files, the intensities of all 11 TMT reporter ions were extracted using MASIC software (v3.2.7465) [45] with the default parameter settings, except for ‘*ReporterIonStatsEnabled value = True*’ and ‘*ReporterIonMassMode value = 16*’ (11-plex TMT). Subsequently, the peptide-spectrum matches (PSMs) of interest were associated with the extracted reporter ion intensities based on the scan number. Signal intensities from tumor and normal samples were segregated according to the TMT reporter ion intensity patterns and the downloaded metadata. Peptide mass spectra was visualized using the *plotSpectra()* function of R package Spectra (v1.5.1).

### Data visualization

Graphical illustrations were generated using the BioRender web tool. Data visualization was performed using R (v4.1.2) and the ggplot2 package (v3.3.6). All figures were combined, readjusted, and rendered in Affinity Designer (Serif, v1.10.5).

### Statistics

For two-group comparisons, Student’s t-test was used unless otherwise specified in figure legends. A significance level of *P* < 0.05 was considered statistically significant. All statistical analyses were performed in R (v4.1.2).

### Study approval

For all human tissue studies, written informed consent was obtained from all patients, and tissue samples were used in accordance with the University of California, San Francisco (UCSF) institutional review board (IRB) for human research. All the experiments and analyses using clinical samples were conducted according to the Declaration of Helsinki.

## Declarations

### Consent for publication

Not applicable.

### Availability of data and materials

Nanopore full-length transcript amplicon sequencing dataset generated in this study will be made available through the NCBI Gene Expression Omnibus (GEO) website upon manuscript acceptance. RNA-seq data of the TCGA (phs000178.v10.p8) and the GTEx (phs000424.v9.p2) datasets were obtained through the GDC Data Portal with an approved dbGaP data access. RNA-seq data of Mayo Clinic GBM PDX is available through the NCBI GEO (accession, PRJNA548556). The CPTAC-GBM proteomics data is available through NIH Proteomic Data Commons (Study ID PDC000204). All requests for materials and correspondence may be sent to.

### Author contributions

Project administration: T.N., H.O.; Conceptualization: T.N., L.W., J.F.C., A.A.D, and H.O; Funding acquisition: T.N., J.F.C., A.A.D, and H.O.; Methodology: T.N., L.W., K.K.L., D.A.C., M.Y.Z., N.O.S., A.Y., P.W., A.N., A.P.W, J.A.W., J.F.C., A.A.D., and H.O.; Data analysis: T.N., L.W., K.K.L., L.H.C., J.F.C., A.A.D., and H.O.; Investigations: T.N., L.W., K.K.L., D.W.K., D.A.C., M.Y.Z., A.Y., P.W., A.N., A.P.W, and H.O.; Resources: A.W., C.H., L.H.C., A.S., J.J.P., S.M.C., and J.F.C.; Supervision: J.A.W., J.F.C., A.A.D., and H.O.; Visualization: T.N.; Writing – original draft: T.N.; Writing – review and editing: S.L., M.G., G.C.M, A.Y., J.F.C., A.A.D., and H.O.; Approval of the final manuscript: all authors.

### Prior publications

Part of this work was presented at the 26^th^ Annual Meeting of the Society for Neuro-Oncology (SNO, 11/20/2021, Boston, MA, USA), and the 6^th^ Quadrennial Meeting of the World Federation of Neuro-Oncology Societies (WFNOS, 03/26/2022, Seoul, Korea).

## Acknowledgments

We thank the following individuals and organizations: the study participants and their families; C4, a shared high-performance compute (HPC) cluster supported by the Helen Diller Family Comprehensive Cancer Center (HDFCCC) and the Computational Biology and Informatics (CBI) Shared Resource, which significantly facilitated the computational analyses conducted in this study; the UCSF Brain Tumor Center SPORE Biorepository and Pathology Core for their services (NIH/NCI 5P50CA097257-18); members of the laboratories of A.P.W., J.A.W., J.F.C., A.A.D., and H.O. for assistance, advice, and helpful discussions.

## Data availability

Nanopore full-length transcript amplicon sequencing dataset generated in this study will be made available through the NCBI Gene Expression Omnibus (GEO) website upon manuscript acceptance. RNA-seq data of the TCGA (phs000178.v10.p8) and the GTEx (phs000424.v9.p2) datasets were obtained through the GDC Data Portal with an approved dbGaP data access. RNA-seq data of Mayo Clinic GBM PDX was obtained through the NCBI GEO (accession, PRJNA548556). The CPTAC-GBM proteomics data was obtained through NIH Proteomic Data Commons (Study ID PDC000204).

## Supplementary Information

**Supplementary Fig. S1.**
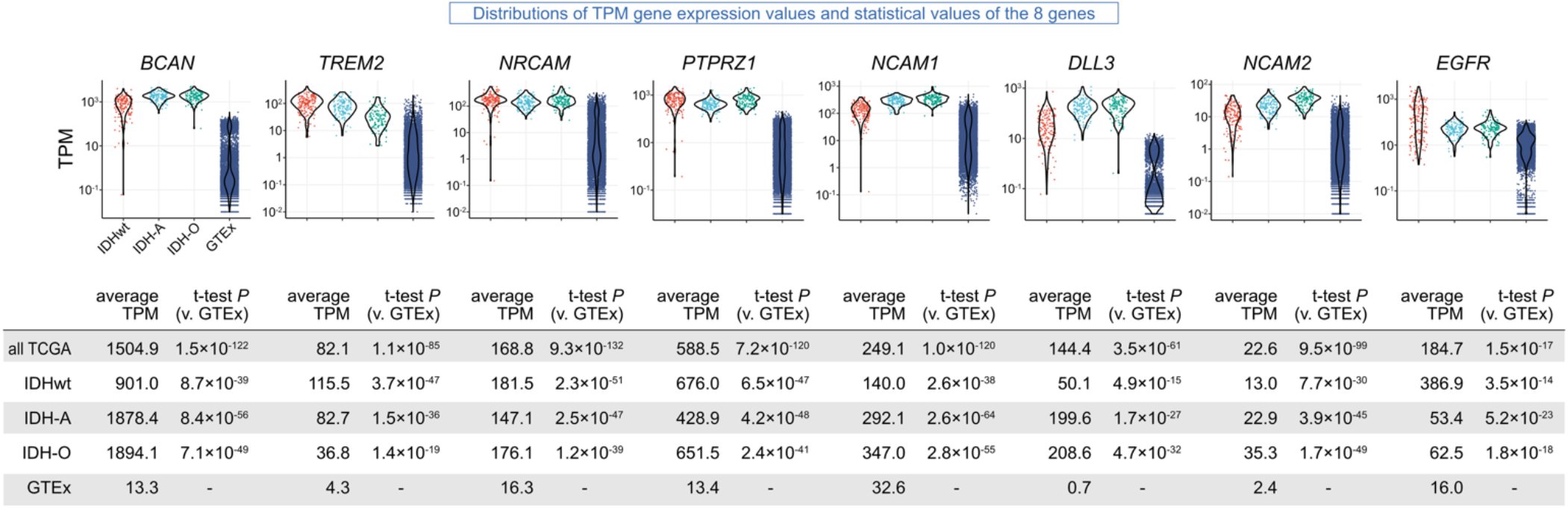
Distribution of gene expression TPM values of the eight genes in the TCGA and the GTEx datasets. Dot and violin plots depicting the gene-level expression of the eight genes that generate the pass-filter candidate AS events in each sample group, identical to Fig. 2b. Average TPM values for all TCGA-glioma, IDHwt, IDH-A, IDH-O, and GTEx groups are displayed in the bottom table, along with corresponding *P*-values calculated using a t-test against the GTEx samples.

**Supplementary Fig. S2.**
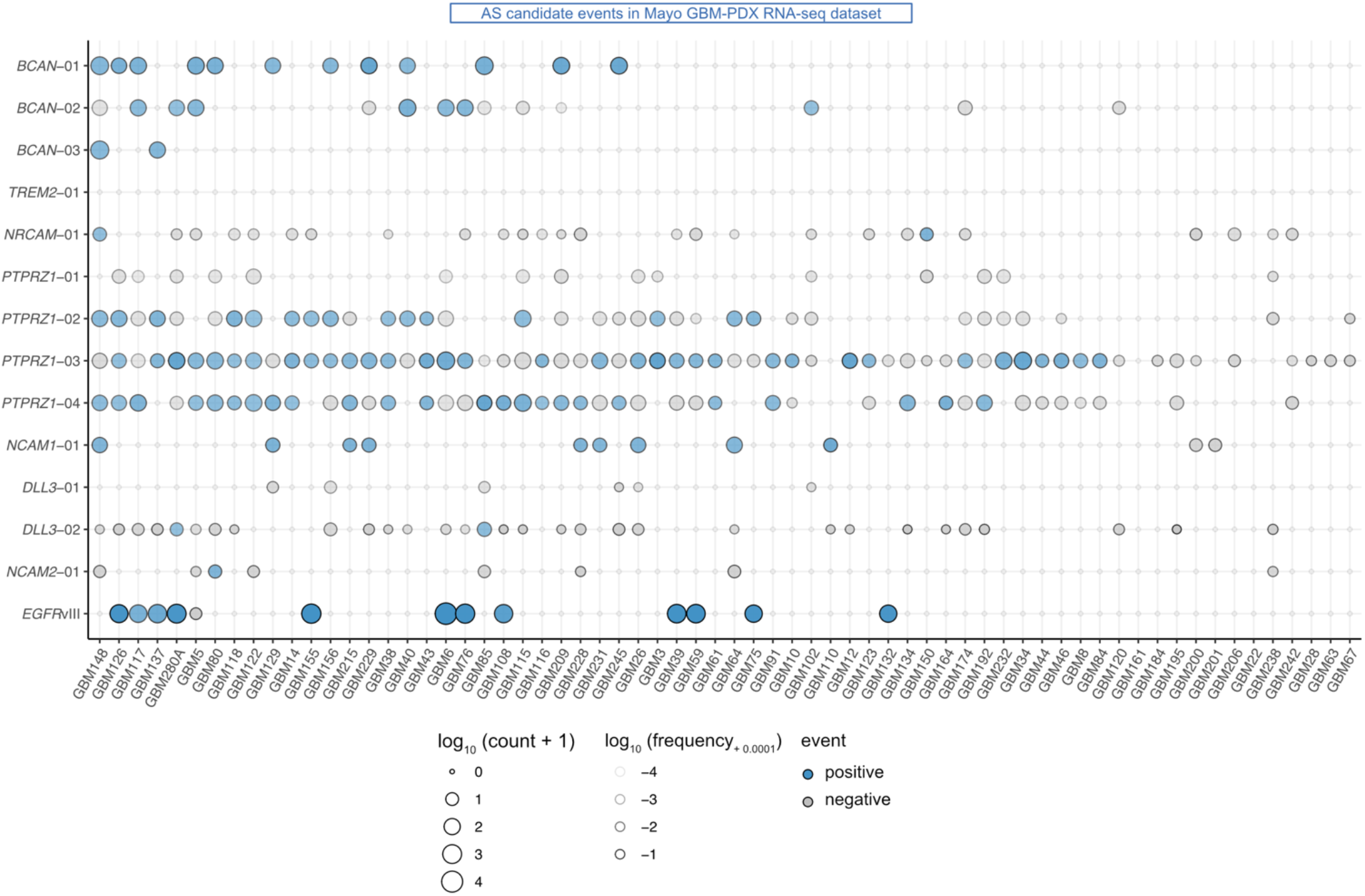
External dataset validation of the identified 13 AS events. Dot plots displaying presence of candidate AS events in the Mayo PDX-GBM transcriptome dataset (n = 66 samples). Blue dots indicate events defined as positive, while gray dots indicate those defined as negative in each sample. The size and color intensity of each dot represent the positive read counts and positive read frequencies, respectively.

**Supplementary Fig. S3.**
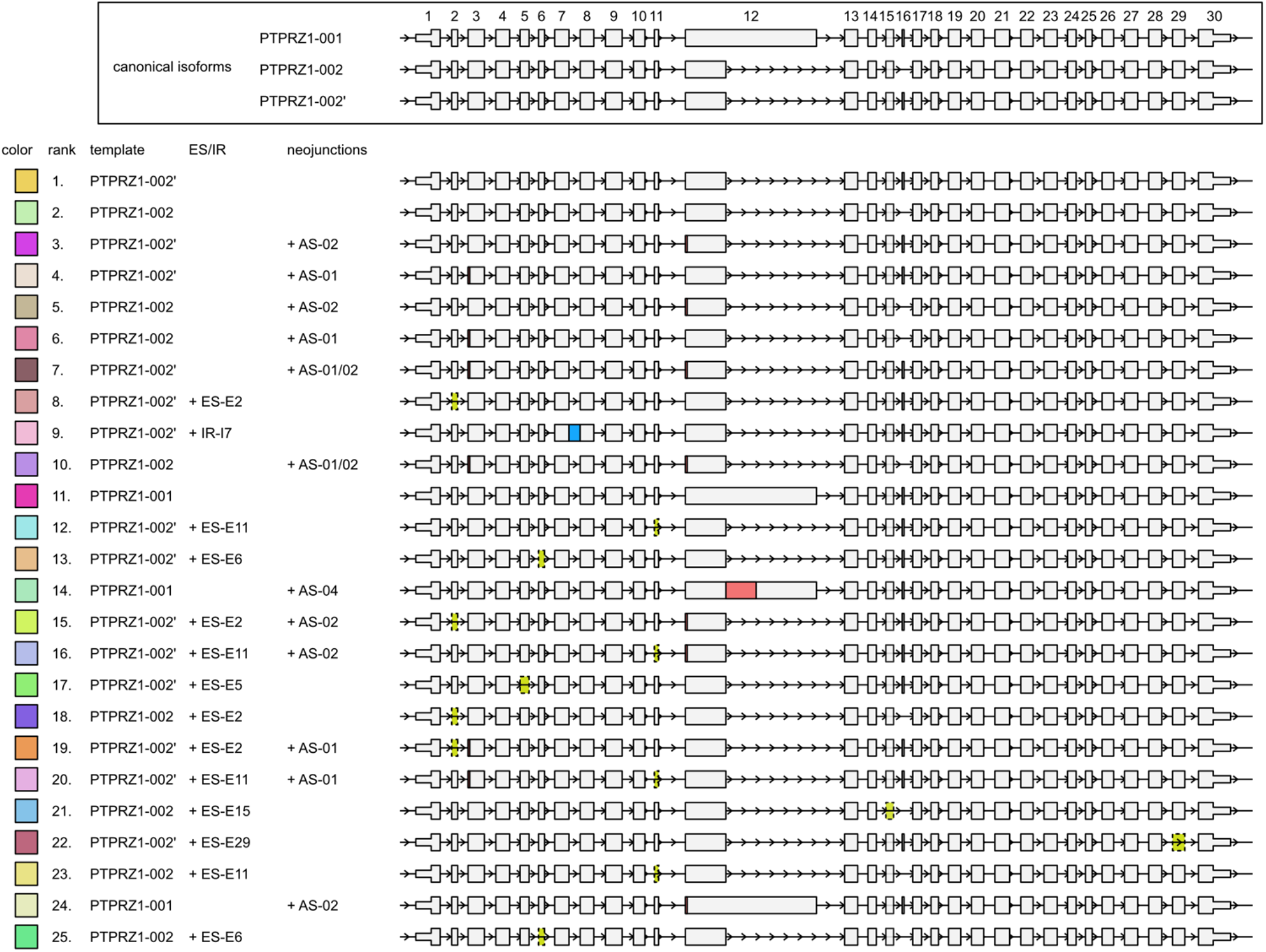
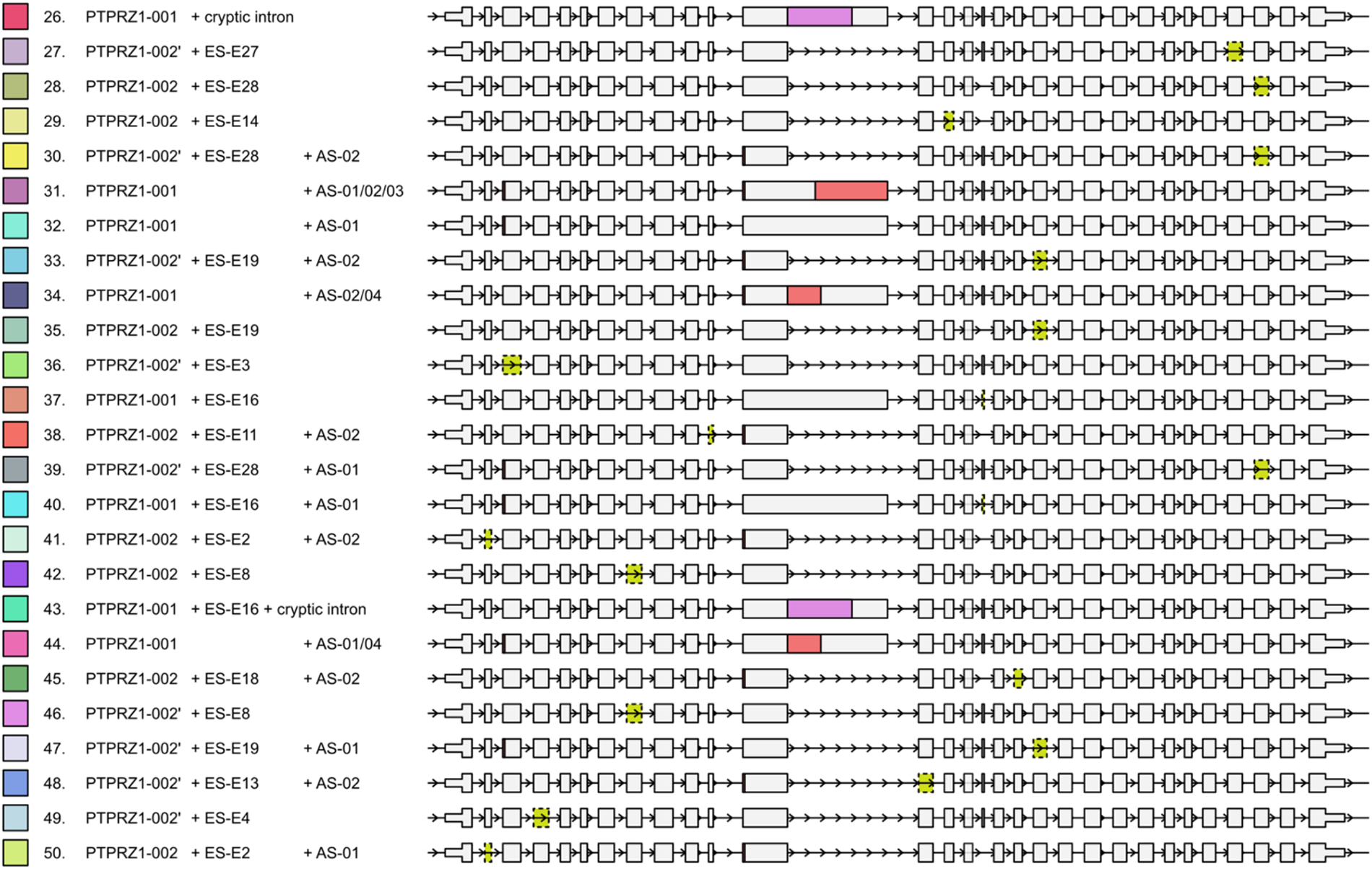
Top 50 isoforms identified in the full-length transcript amplicon sequencing. Schematic illustration of the top 50 most frequently identified isoforms including canonical and cryptic. The colors in the left square correspond to Fig. 3e. The rank number indicates the ranking in the list. Predicted AS patterns (PTPRZ1-01, -02, -03, and -04) (red), and other exon skipping (yellow), intron retention (blue), and cryptic intron (purple) events are highlighted.

**Supplementary Fig. S4.**
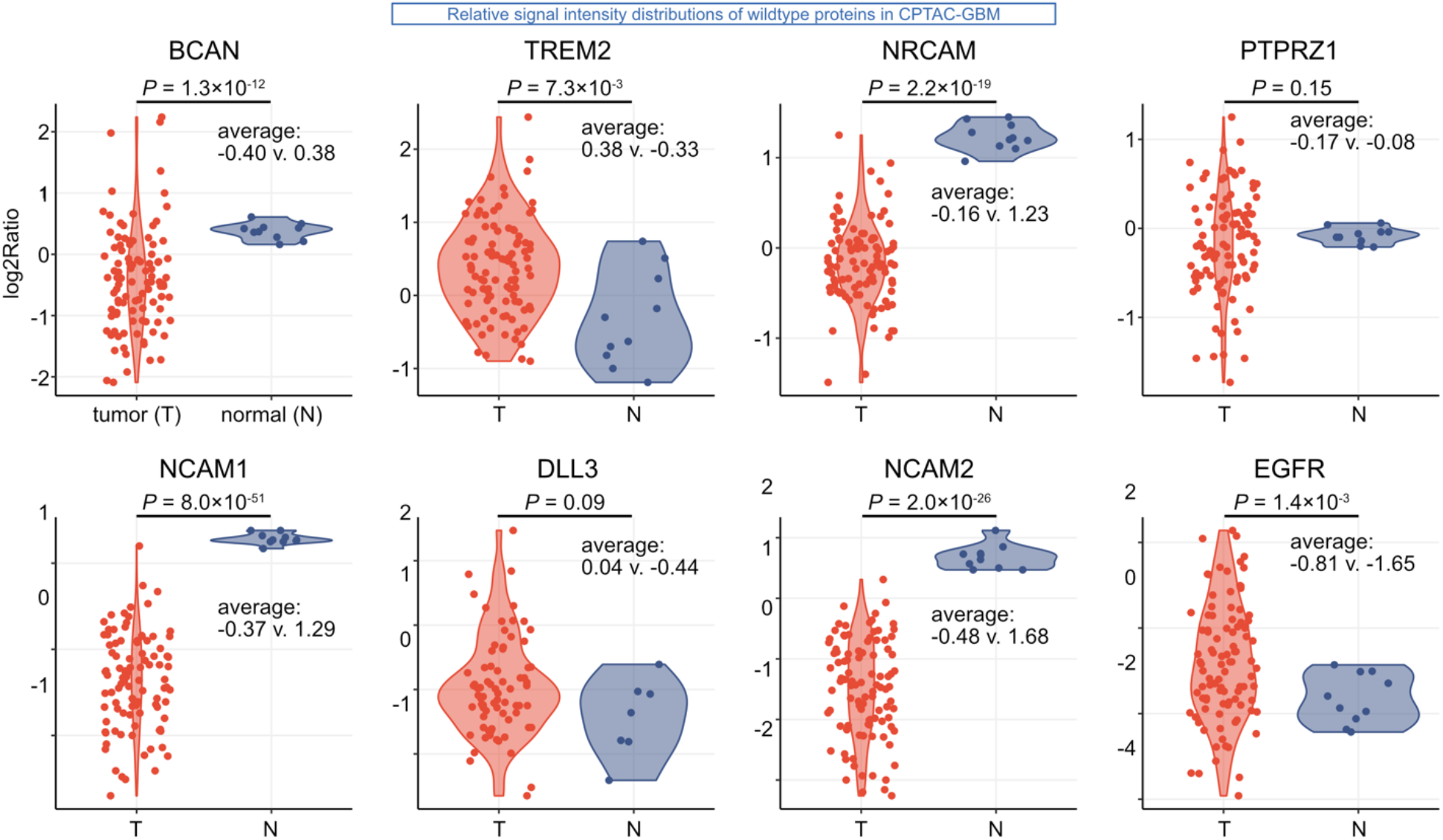
Relative signal intensity distributions of wildtype proteins corresponding to the candidate AS events. Dot and violin plots showing the distributions of relative signal intensities for each of the wild-type proteins corresponding to the candidate AS events. The data has been sourced from the CPTAC-GBM data portal and is presented in its raw form, without additional modifications or transformations. The Y-axis represents log2 ratios to internal control reference samples.

**Supplementary Fig. S5.**
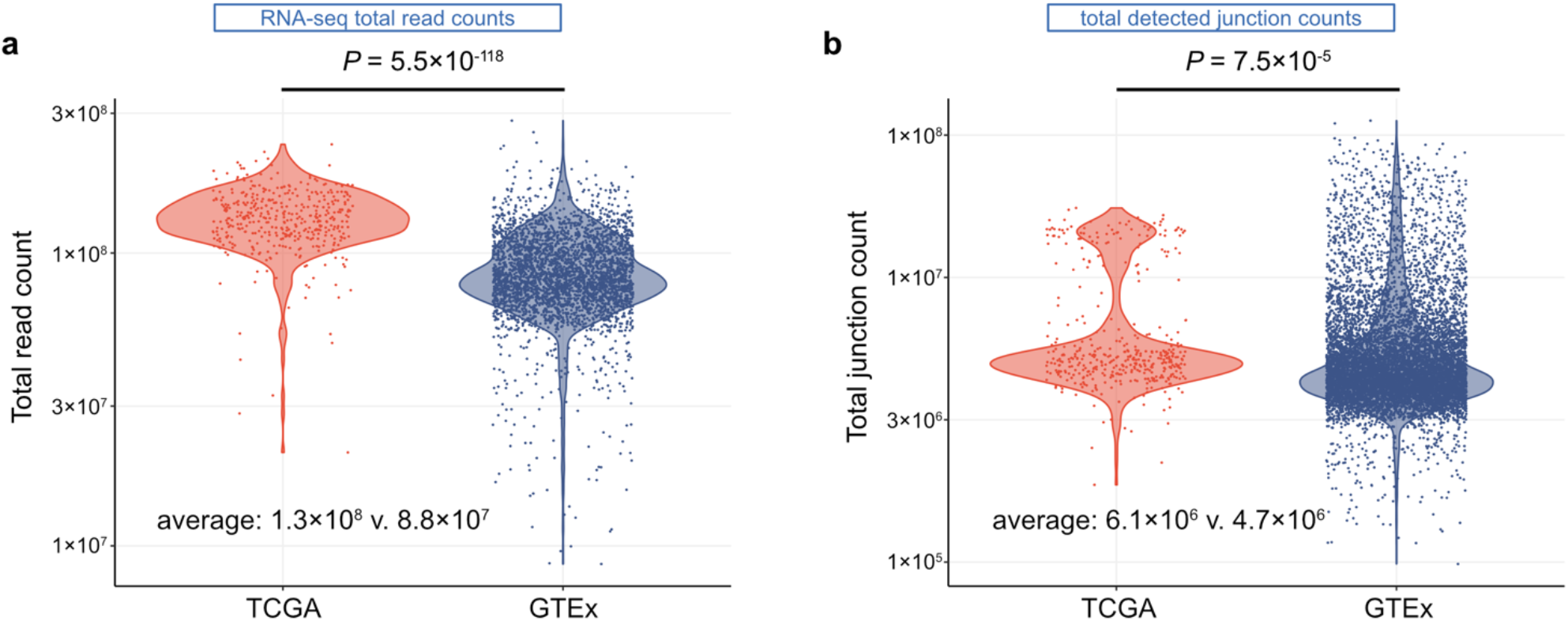
Distributions of RNA-seq total read counts and detected junction counts between the TCGA and the GTEx datasets. Dot and violin plots showing the distributions of total read counts within the RNA-seq BAM file (**a**) and the total unique junction counts within the SJ.out.tab file (**b**) of each sample. *P* values are calculated using *t-*test.

